# A Subset of Olfactory Sensory Neurons Express Forkhead Box J1-Driven eGFP

**DOI:** 10.1101/643460

**Authors:** Shivani Pathak, Eric D. Larson, Vijay R. Ramakrishnan, Thomas E Finger

## Abstract

Forkhead box protein J1 (*Foxj1*), a member of the forkhead family transcription factors, is a transcriptional regulator of motile ciliogenesis. The nasal respiratory epithelium, but not olfactory epithelium, is lined with FOXJ1-expressing multiciliated epithelial cells with motile cilia. Using a *Foxj1*-eGFP reporter mouse, we find robust eGFP expression not only in the multi-ciliated cells of the respiratory epithelium, but in a distinctive small subset of olfactory sensory neurons in the olfactory epithelium. These eGFP-positive cells lie at the extreme apical part of the neuronal layer and are most numerous in dorsal-medial regions of olfactory epithelium. Interestingly, we observed a corresponding small number of glomeruli in the olfactory bulb wherein eGFP-labeled axons terminate, suggesting that the population of eGFP+ receptor cells expresses a limited number of olfactory receptors. Similarly, a subset of vomeronasal sensory neurons express eGFP but these distribute throughout the full height of the vomeronasal sensory epithelium. In keeping with this broad distribution of labeled vomeronasal receptor cells, eGFP labeled axons terminate in many glomeruli of the accessory olfactory bulb. These findings suggest that *Foxj1*-driven eGFP marks a specific population of olfactory and vomeronasal sensory neurons although neither receptor cell population possess motile cilia.

## Introduction

The nasal cavities of mice are lined by two distinct epithelial surfaces: the respiratory epithelium (RE) situated anteriorly and the olfactory epithelium (OE) situated posteriorly. A key feature of the respiratory epithelium is the multiciliated cell with motile cilia. The OE is a pseudostratified epithelium predominantly comprising of olfactory sensory neurons (OSNs) lying below a single apical layer of sustentacular cells. OSNs contain the necessary cellular machinery for detection of odorants and transmission of related neuronal signals to the olfactory bulb. OSN dendrites project apically towards the nasal cavity and terminate in a knob at the epithelial surface from which multiple non-motile sensory cilia extend.

The main OE is mapped into four zones -- arranged dorso-medial to ventro-lateral -- based on patterns of odorant receptor (OR) expression. Each mature OSN expresses only one predominant OR and is restricted to an individual zone. The axons of these OSNs then converge onto one or two glomeruli within the main olfactory bulb (MOB), such that each glomerulus receives input from receptor cells expressing only one OR (Hanchate et al., 2015; Mori, von Campenhause, & Yoshihara, 2000).

The olfactory system of mice also includes the vomeronasal organ (VNO), a sensory organ that lies along either side of the floor of the main nasal cavity. An anterior duct directs fluids into the lumen of the organ posteriorly where the vomeronasal sensory neurons (VSNs) are located. This epithelium is also organized by chemoreceptor expression, but, instead of 4 zones, the SE is subdivided into 2 layers of cell bodies, apical and basal, that respectively express the V1R and V2R classes of vomeronasal receptors. These neurons relay information through the vomeronasal nerve located within the septum to the Accessory Olfactory Bulb (AOB). Specifically, axons of the apical neurons project to the anterior accessory olfactory bulb (AOB) while axons of the basal neurons project to the posterior AOB (Breer, Fleischer, & Strotmann, 2006). Interestingly, receptor neurons in the VNO do not possess cilia but rather actin-based microvilli (Doving & Trotier, 1998; Stensaas, Lavker, Monti-Bloch, Grosser, & Berliner, 1991; Yoshida-Matsuoka, Osada, Mori, & Ichikawa, 1999).

In this study, we use a transgenic mouse model developed by Ostrowski and colleagues to study respiratory cilia, in which eGFP expression is linked to the *Foxj1* promoter(Ostrowski, Hutchins, Zakel, & O’Neal, 2003), a transcription factor required for motile ciliogenesis for mono and multiciliated cells (You et al., 2004; Yu, Ng, Habacher, & Roy, 2008). As expected, we observed eGFP expression within the RE (Figure 1), since the predominant cell type of this epithelium is the multiciliated cell and *Foxj1* is a known driver of motile ciliogenesis (Blatt, Yan, Wuerffel, Hamilos, & Brody, 1999; Hackett et al., 1995; Lim, Zhou, & Costa, 1997). Surprisingly, we observed expression of eGFP in sub-populations of OSNs and VSNs neither of which possess motile cilia. Here, we further describe this population of OSNs including their distribution within the main OE and VNO and the concomitant projections of their axons to the MOB and AOB.

**Figure 1.**
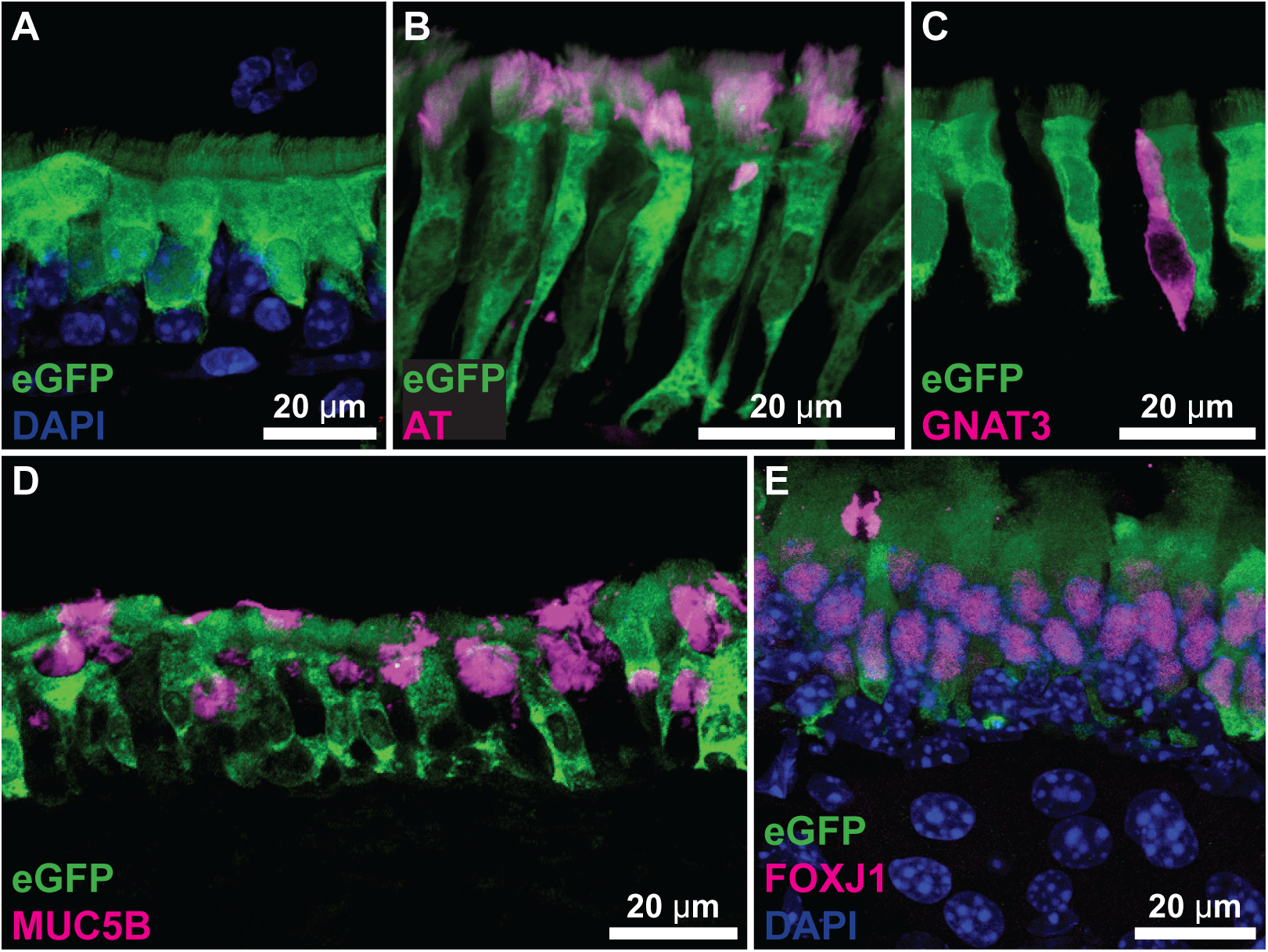
Respiratory Epithelial (RE) Staining. Foxj1-driven eGFP (green) robustly labels multi-ciliated epithelial cells. A-B. Multi-ciliated epithelial cells of the nasal RE express Foxj1-driven EGFP throughout the cell body and cilia marked by acetylated tubulin (magenta). C. Solitary chemosensory cells identified by staining for GNAT3, do not express GFP. D. Goblet cells, visualized by MUC5B immunoreactivity are distinct from eGFP-expressing multi-ciliated cells. E. FOXJ1 antibody (magenta) reveals protein is present within the nuclei of GFP-labled multi-ciliated cells. Images are maximum z-projections of 5-10 µm substacks.

## Materials and Methods

### Animals

All use of experimental animals was approved by the Institutional Animal Care and Use Committee and the University of Colorado Anschutz Medical Campus (Aurora, CO). All experiments were conducted on male and female mice aged 2-6 months, housed in ventilated cages on a 14-hr light cycle with *ad libitum* access to standard chow. The *Foxj1*-eGFP mouse line (B6;C3-Tg(FOXJ1-EGFP)85Leo/J) generated by Ostrowski et. al. (RRID:IMSR_JAX:010827) has been described previously (Ostrowski et al., 2003).

### Anesthetic and Fixation

All mice were sacrificed at 2-6 months and were either immersion-fixed overnight, or transcardially perfused using 4% paraformaldehyde in 0.1M phosphate buffer (PB). There were no appreciable differences between sexes or method of fixation. Dissection involved removing parts of the skull to expose the olfactory bulbs, forebrain, and nasal cavities. The head was decalcified using 0.45 M EDTA (pH 8) for 24-36 hours. All heads underwent cryoprotection overnight with 20% sucrose in 0.1M PB. Tissue was then embedded in OCT compound (Optimal Cutting Temperature; Fisher Scientific, Pittsburgh, PA) and cut on a cryostat. 16-micron sections were collected directly onto charged glass microscope slides (Light Labs USA, Dallas, TX).

### Immunohistochemistry and Imaging

After one 10-minute wash in 0.1M PB and two 10-minute washes in PBS (phosphate buffered saline, pH 7.4), all slides underwent antigen retrieval using 10mM sodium citrate (pH 9) buffer at 85 °C for 25 minutes. After cooling, two additional PBS washes were performed prior to incubating with blocking solution (2% normal donkey serum, 1% bovine serum albumin, 0.3% triton) for 1 hour. Primary antibodies were diluted in blocking solution, applied to the slides and incubated overnight at 4°C. For a complete list of antibodies used in this study, refer to Table 1. The following day, slides were washed in PBS solution three times and then incubated with the appropriate secondary antibodies (Table 1) for 3 hours. After 3 washes, slides were counterstained with DAPI and mounted using Fluoromount-G (Southern Biotech). All sections were viewed with an epifluorescence microscope and imaged on a Leica SP5 or SP8 laser scanning confocal microscope equipped with 20x (NA 0.75) and 63x (NA 1.4) objectives. **Characterization of Primary Antibodies (see Table 1)**

**Table 1.** Primary and Secondary Antibodies.

| **Antiserum** | **Company** | **Catalog**<br>**Number** | **Dilution** | **RRID** |
| --- | --- | --- | --- | --- |
| Chicken Polyclonal Anti-GFP | Aves Lab; Trigard, OR | NC9510598 | 1:2000 | AB_1000024<br>0 |
| Rabbit polyclonal anti-OMP | Sigma; St. Louis, MO | MFCD09265364 | 1:500 | AB_796160 |
| Goat polyclonal anti-GNAT3 | Aviva Systems Bio; San<br>Diego, CA | OAEB00418 | 1:500 | AB_1088282<br>3 |
| Mouse Monoclonal anti FOXJ1 | Thermo Fisher, Waltham,<br>MA | 14-9965-82 | 1:500 | AB_1548835 |
| DAPI | Invitrogen; Carlsbad, CA | 62248 | 1:10,000 | AB_2307445 |
| Donkey anti-Rabbit 568 | Invitrogen; Carlsbad, CA | A10042 | 1:800 | AB_2534017 |
| Donkey anti-Chicken 488 | Jackson Immuno Research;<br>West Grove PA | 703-546-155 | 1:800 | AB_2340375 |
| Donkey anti-Goat 647 | Invitrogen; Carlsbad, CA | A-21447 | 1:800 | AB_2535864 |
| DRAQ5™ | Abcam; San Francisco, CA | AB_108410 | 1:5000 | AB_108410 |
| Mouse anti-GFAP | Sigma; St. Louis, MO | SAB1405864 | 1:300 | AB_477010 |
| Rabbit Polyclonal anti-MUC5B | Santa Cruz Bio; Santa Cruz,<br>CA | sc-20119 | 1:200 | AB_1842559 |
| Mouse anti-Acetylated Tubulin | Sigma; St. Louis, MO | **T7451** | 1:2000 | AB_609894 |

#### Green Fluorescent Protein (GFP; Aves)

This antibody recognizes the full length green fluorescent protein by Western blot analysis and explicitly co-labels GFP in GFP-expressing transgenic mice (Aves datasheet). We observe no labeling of wildtype mice lacking transgenic GFP expression.

#### Olfactory Marker Protein (OMP; Sigma)

This rabbit polyclonal antibody recognizes amino acids 119-137 near the C-terminus of rat OMP and based on sequence homology is predicted to recognize mouse OMP. Analysis by Western blot shows a band at 19 kDA from rat olfactory bulb which is blocked by inhibition with specific OMP immunizing peptide (Sigma datasheet).

#### Guanine nucleotide-binding protein G(t) subunit α-3 (GNAT3; Aviva)

This polyclonal antibody produced in goats recognizes the epitope ‘C-KNQFLDLNLKKEDKE’ from an internal region of human GNAT3 protein. Based on sequence homology it is predicted to recognize mouse, rat, rabbit, pig, dog, and cow GNAT3. Western blot analysis shows specific recognition of a 27 kDa band from human colon tissue (Aviva datasheet). Further, we observe no signal when tested on nasal section from animals lacking the transcription factor *Pou2f3* (Yamashita, Ohmoto, Yamaguchi, Matsumoto, & Hirota, 2017) which is required for the generation of solitary chemosensory cells.

#### FOXJ1 (FOXJ1; Thermo Fisher)

This mouse monoclonal antibody reacts with human and mouse FOXJ1. Analysis by Western blot shows a 60 kDa band in mouse tracheal epithelial cells. Additionally, FOXJ1 was detected by Western blot in human bronchial epithelial cells before but not after siRNA knockdown of *Foxj1* (Jacquet et al., 2009).

#### Mucin5B (MUC5B; Santa Cruz)

This rabbit polyclonal antibody recognizes an epitope within amino acids 1201-1500 of human Mucin5B and is predicted to recognize mouse Mucin5B. Staining pattern of this antibody is consistent with goblet cell expression (Figure 1).

#### Acetylated Tubulin (AT; Sigma)

This mouse monoclonal antibody reacts with multiple species including mouse. Per company datasheet, the antibody reacts with a region of α3 isoform of *Chlamydomonas* axonemal α-tubulin and analysis by Western Blot shows a ∼55 kDa band in lysates from multiple cell lines. Staining patterns are consistent with microtubules.

#### Glial Fibrillary Acidic Protein (GFAP; Sigma)

This mouse monoclonal antibody was produced against the full length human protein GFAP. Analysis by Western Blot shows a band of approximately 50 kDa.

### Image quantification

All quantification was performed using the FIJI distribution of ImageJ (v1.52n; (Schindelin et al., 2012)). 20× tile-scan images were background subtracted (‘Subtract Background’, rolling ball radius of 50 px) across all channels. ROIs were drawn around individual glomeruli as identified by OMP fluorescence for measurement of mean fluorescence intensity. For colocalization analysis, high-magnification images were processed as above. Colocalization within specific glomeruli was measured using Coloc2 plugin bundled with FIJI. Plots were made in R ((2008).)with *ggplot2* (Wickham., 2016)

## Results

As expected, multiciliated cells of the RE have robust cytoplasmic eGFP expression consistent with *Foxj1’s* known role as a driver of motile ciliogenesis. However, we noted expression of eGFP in a sub-population of OSNs and in receptor cells of the VNO, neither of which have motile cilia. To further characterize expression in these systems, we analyzed the distribution of eGFP-label in the mouse olfactory system including the main olfactory epithelium, the main olfactory bulb (MOB), the vomeronasal organ (VNO), and the accessory olfactory bulb (AOB).

### Respiratory epithelium

Multiciliated cells of the respiratory epithelium have robust eGFP expression (Figure 1) extending into the cilia of these cells, co-labeled by acetylated tubulin (Figure 1B). We did not detect eGFP expression in the respiratory epithelium in cells lacking motile cilia: solitary chemosensory cells (GNAT3-expressing; Figure 1C) or goblet cells (MUC5B-expressing; Figure 1D). The nuclei of the eGFP-labeled multiciliated cells are immunoreactive for FOXJ1 (Figure 1E) indicating that FOXJ1 protein is present in these cells consistent with the presence of *FoxJ1*-driven eGFP. These findings are consistent with the known function of *Foxj1* driving motile ciliogenesis (Blatt et al., 1999; Hackett et al., 1995; Lim et al., 1997).

### Main olfactory epithelium (MOE)

The olfactory epithelium of *Foxj1*-eGFP mice show a subset of OSNs that co-express *Foxj1*-driven eGFP and OMP (Figure 2). eGFP immunoreactivity within OSNs is robust and often can be visualized in the immotile cilia and axons (Figure 2B, C). All neurons expressing eGFP also express OMP, confirming that they are, in fact, mature OSN cell bodies. Further, the cell bodies of these neurons are consistently situated apically within the OMP positive layer of the epithelium. Since OR expression is laminarly segregated (Strotmann, Konzelmann, & Breer, 1996), the consistent apical situation of the eGFP-labeled OSNs suggests that the eGFP expression may correlate with expression of particular ORs. In particular, the apical situation of the eGFP-labeled OSNs is similar to the location of members of the mOR37 family as reported by Strotman and co-workers (Strotmann et al., 2000; Strotmann et al., 1996). The *Foxj1*-eGFP-OSNs preferentially populate the dorsomedial region of the MOE (Figure 3), suggesting these neurons are zonally organized, similar to the zonal expression of ORs across the MOE (Hanchate et al., 2015; Mori, Takahashi, Igarashi, & Yamaguchi, 2006). Taken together, these findings suggest that *Foxj1*-eGFP-expressing OSNs are mature OSNs that express a specific subset of olfactory receptors. Since olfactory receptor neurons expressing a common receptor project to 1 or 2 glomeruli in the MOB, we examined the MOB to assess whether that the eGFP-labeled OSNs project to a limited number of glomeruli in the MOB.

**Figure 2.**
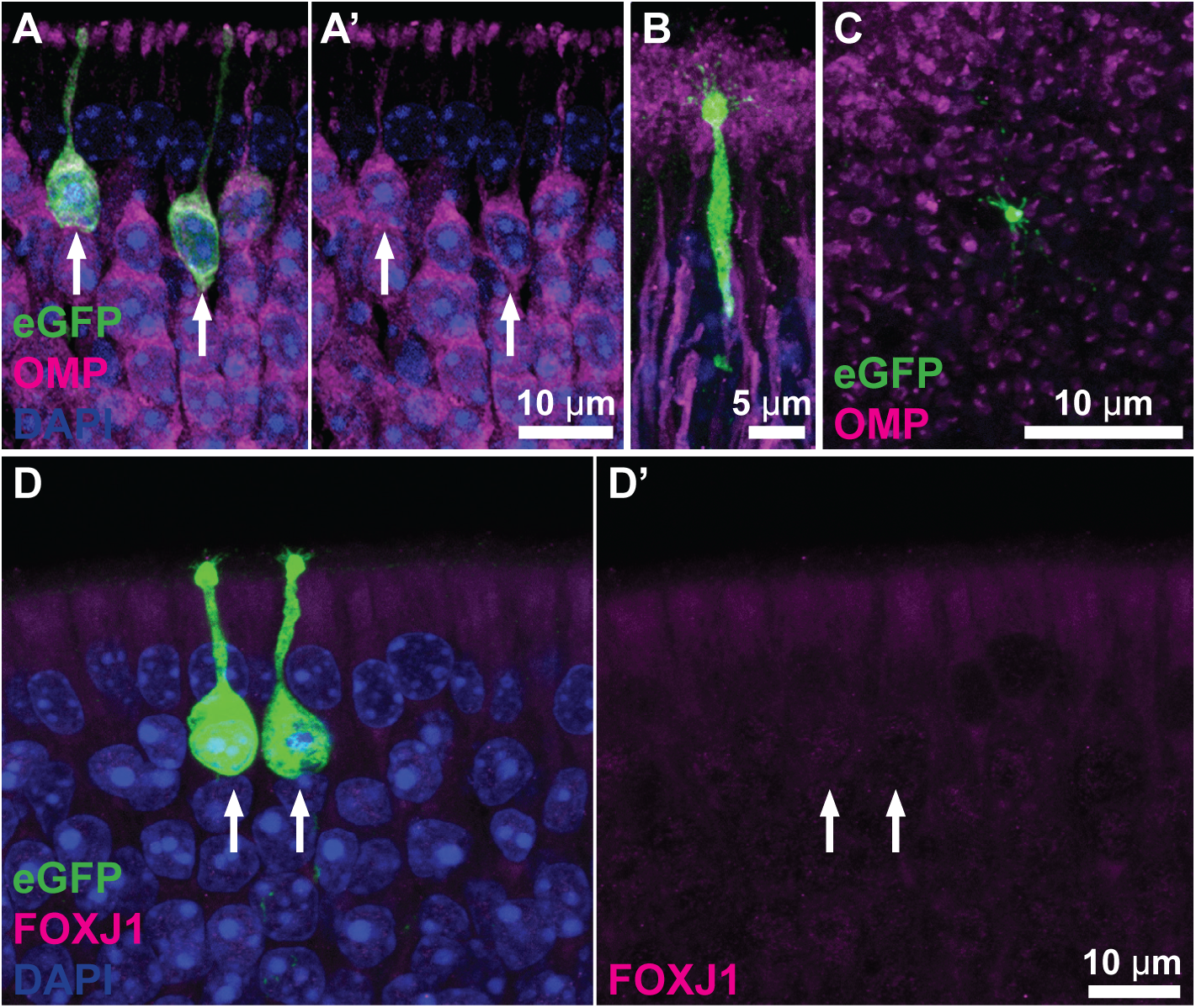
A subset of OSNs express both OMP (magenta) and Foxj1-driven eGFP (green). A. eGFP expressing neuronal cell bodies are located apically within the OE (arrows) and are immunoreactive for OMP. A’. Identical image to panel A but showing only the OMP label (magenta). B, C. eGFP expression in the dendrite, dendritic knob, and cilia of the OSN as viewed in longitudinal section (B) and when the surface of the epithelium is viewed en face (C). D. The nuclei and cell bodies of the eGFP expressing OSNs are not immunoreactive for FOXJ1 (magenta). D’. Identical image to panel D but showing only the FOXJ1 label. Images are maximum z-projections of 5-10 μm substacks.

**Figure 3.**
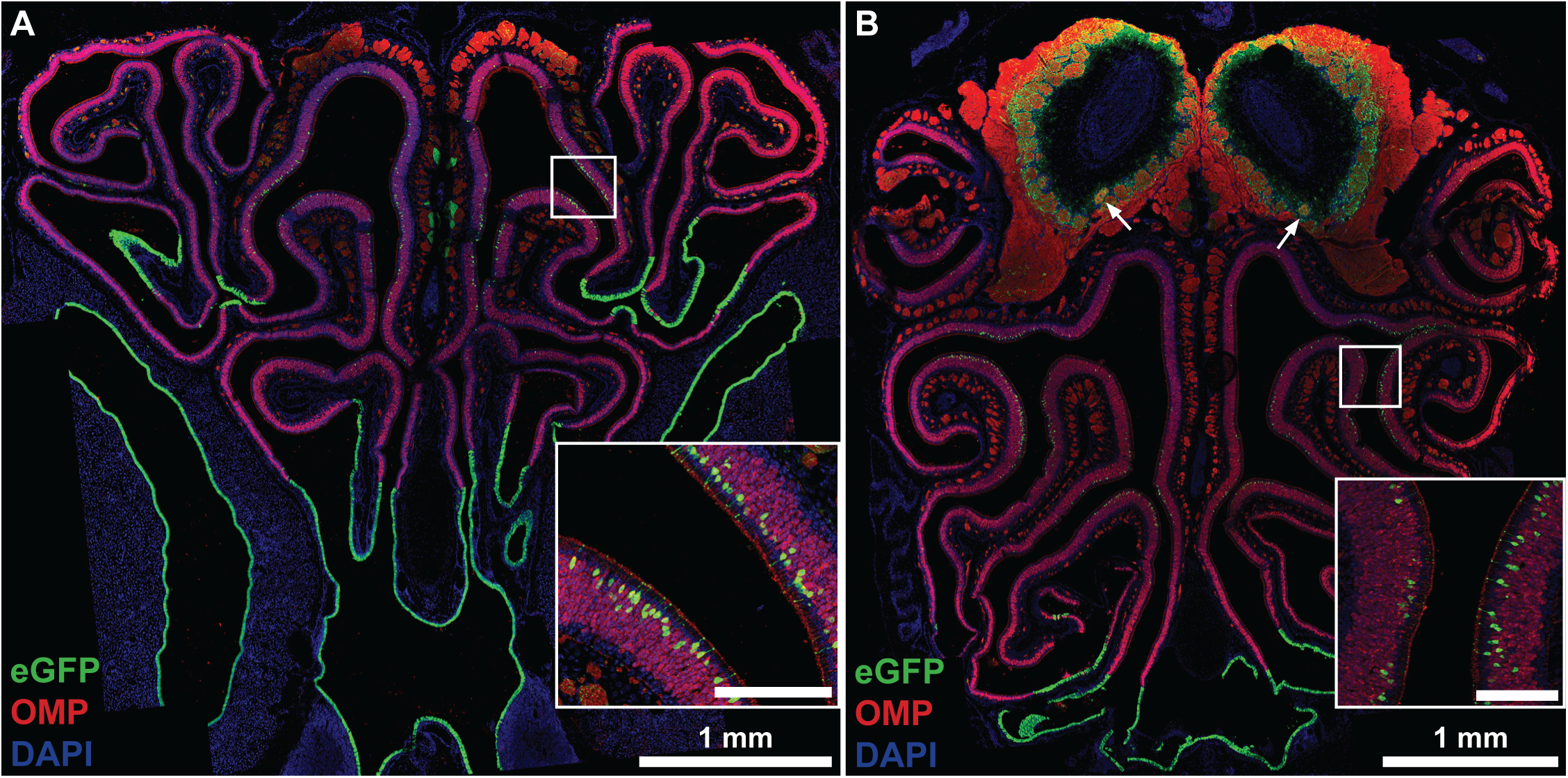
The majority of eGFP expressing OSNs are restricted zonally (dorso-medial) within the MOE. This zonal organization is conserved throughout the MOE as depicted in transverse sections from different regions (A-more anterior, B-more posterior). Two eGFP targeted glomeruli are visible in B (arrows). Insets are magnifications of region denoted by the white box. Inset scale bars, 100 µm. Images are maximum z-projections of a 20× tile-scans.

### Main Olfactory Bulb (MOB)

Numerous branched glial cells throughout the olfactory bulb express eGFP in the FoxJ1-eGFP mouse as reported previously (Jacquet et al., 2009). Some of these GFP-labeled glial cells express GFAP at low levels suggesting that the labeled cells are astrocytes (Figure 4) (Cahoy et al., 2008). The ubiquitous presence of these cells hindered identification of glomeruli receiving projections from the eGFP-labeled OSNs. Nonetheless, double labeling with OMP allowed us to identify glomeruli in which the glomerular neuropil was labeled, thereby identifying glomeruli receiving input from eGFP-labeled OSNs.

**Figure 4.**
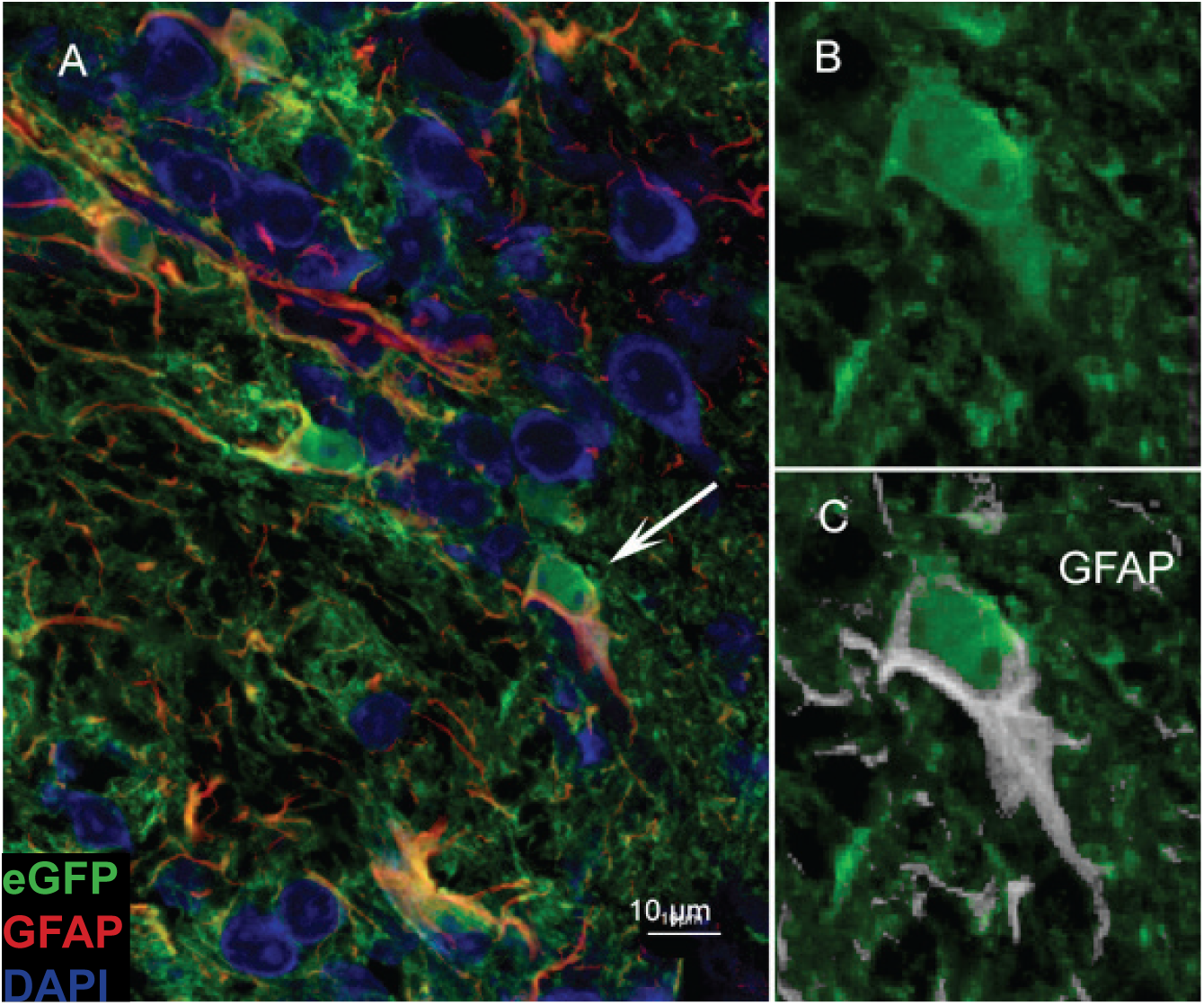
Numerous branched cells in the olfactory bulb expressed GFP. Staining with GFAP antibody (magenta, main image; white inserts at right) confirms their identity as a type of astrocyte as reported previously (Jacquet et al., 2009). A. Double label image of the olfactory bulb showing double label branched cells. Arrow indicates the cell enlarged in panels B & C. B. GFP labeling shows a large multipolar cell. C. Superimposed GFAP label (white) shows label surrounding the nucleus and extending into the proximal extensions of the soma as is typical of GFAP label of astrocytes.

Using this double-label approach, we identified only 5 glomeruli (out of 1631) in 1 hemisphere of a MOB from one animal that had coincident eGFP/OMP fluorescence indicating a convergence of axonal fibers from neurons in the MOE (Figure 5A). These glomeruli were located in the anterior, ventral region of the olfactory bulb and this location was consistent across 4 additional animals (Figure 5B). Quantitative colocalization analysis on 5 labeled and 20 unlabeled glomeruli (Figure 6C) within the ventral anterior MOB further confirmed that eGFP was co-expressed in the OSN axons terminating in a subset of glomeruli (Li’s ICQ value labeled: 0.13 ± 0.02; unlabeled: −0.14 ± 0.008, p<0.0001 by Mann-Whitney U test). As an alternative quantification, we measured the eGFP to OMP immunoreactivity fluorescence ratio in all imaged glomeruli. Indeed, 5 of 1631 glomeruli had a significantly increased ratio (Figure 6D) and were deemed outliers from the dataset by Rosner’s Extreme Studentized Deviate test for multiple outliers (p<0.05) as well as by Iglewicz and Hoaglin’s robust test for multiple outliers (modified Z score≥3.5). This suggests that the eGFP labeled OSNs target specific glomeruli and they are a distinct subset of neurons in regard to OR expression.

**Figure 5.**
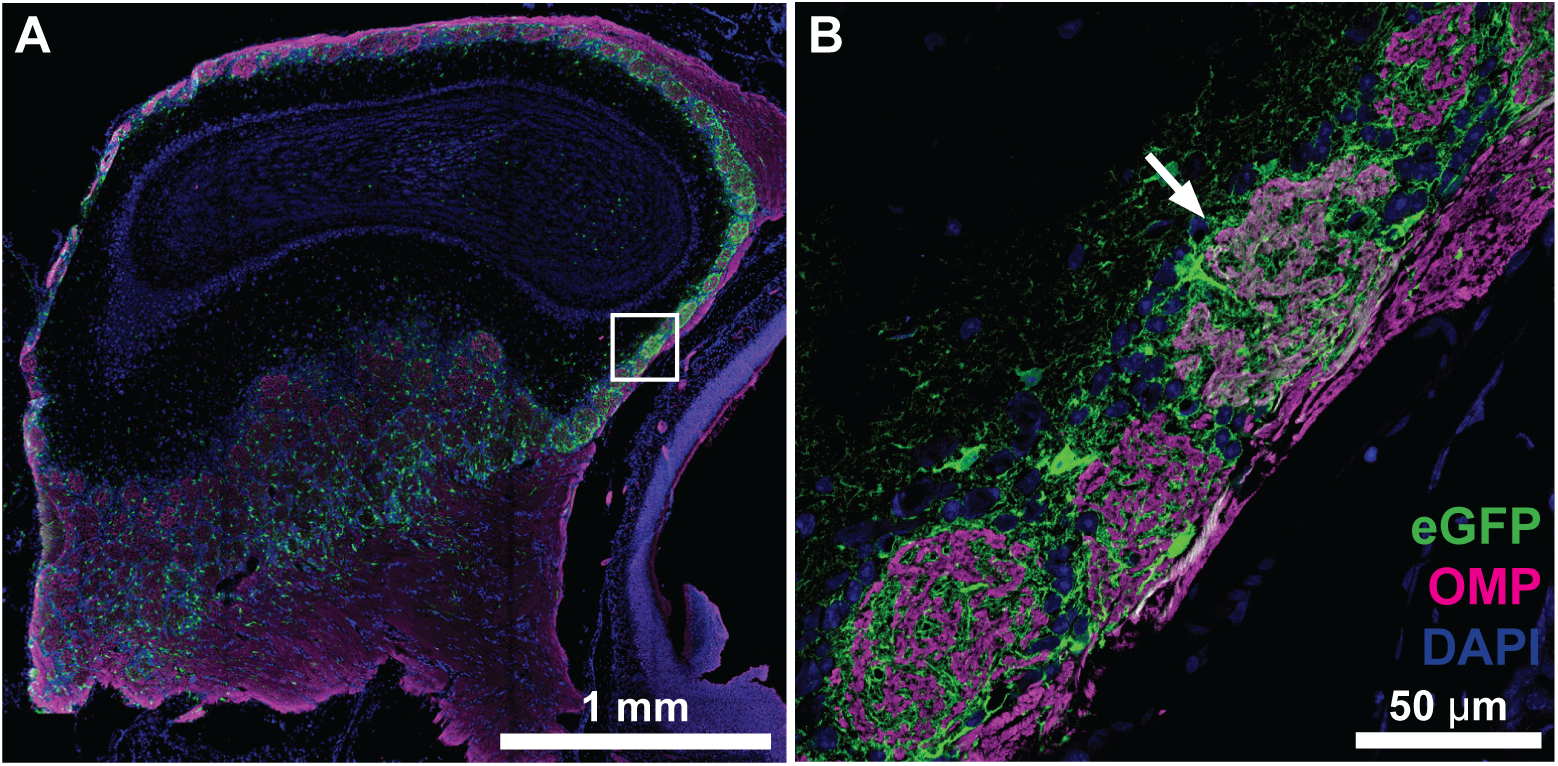
Foxj1-driven eGFP (green) positive neurons converge onto individual glomeruli within the MOB. A. A sagittal 20x tile scan of a section of the MOB shows an individual glomerulus targeted by eGFP (white box). Dorsal is towards the top of the page. B. Higher magnification image of the region depicted in the white box in A shows eGFP neurons converging onto a single glomerulus within the OB. While eGFP-expressing astrocytes are present in all glomeruli, the targeted glomerulus has a higher co-incidence of eGFP and OMP.

**Figure 6.**
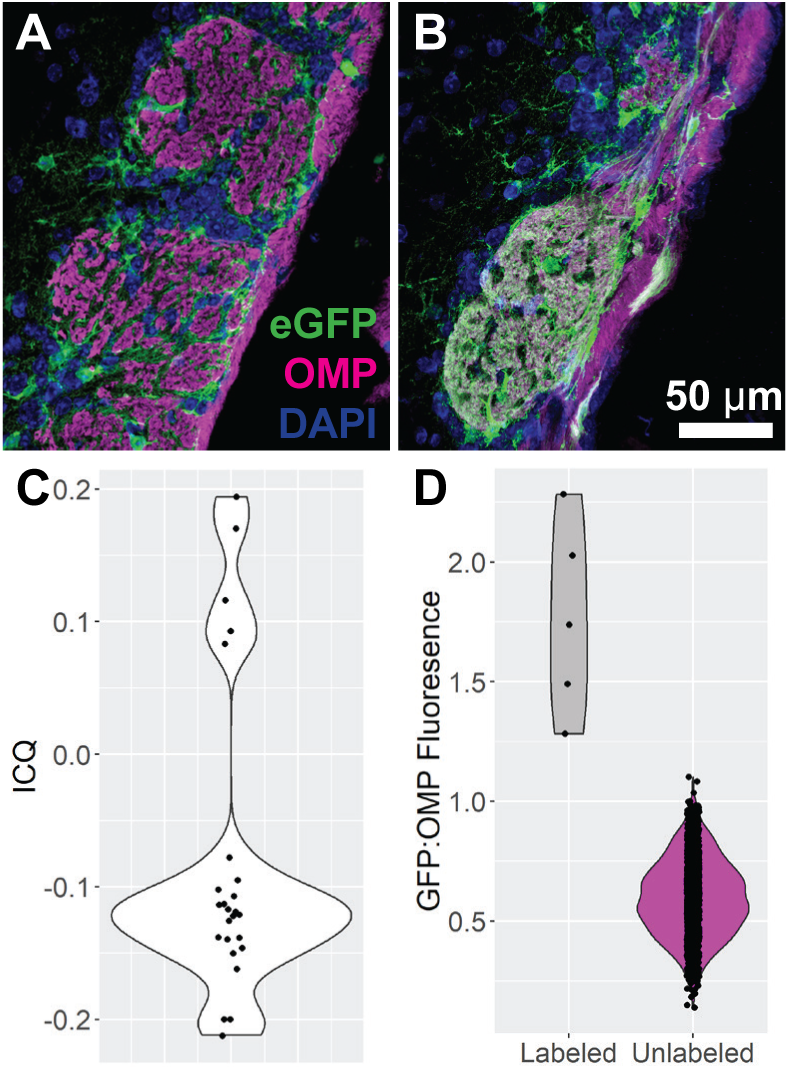
Quantification of glomerular fluorescence intensity reveals eGFP-targeted glomeruli A. Glomerulus in MOB showing neuropil labeled by OMP only. B. Glomerulus in MOB showing neuropil double-labeled for OMP and eGFP. C. Colocalization analysis of 25 glomeruli including the 5 we determined were qualitatively determined as targeted by eGFP-expressing axons. Li’s ICQ value was significantly higher, demonstrating stronger colocalization between the two signals in “targeted” glomeruli. D. Fluorescence intensity was measured in all imaged glomeruli (1631) and the ratio of eGFP to OMP fluorescence was calculated. The 5 glomeruli identified as being targeted by eGFP-expressing axons displayed significantly higher ratios of eGFP:OMP fluorescence and were significant outliers when all datapoints were analyzed by Rosner’s Extreme Studentized Deviate test for multiple outliers as well as by Iglewicz and Hoaglin’s robust test for multiple outliers.

### Accessory Olfactory System - Vomeronasal Organ and Accessory Olfactory Bulb

Within the VNO, we see scattered expression of eGFP in both the apical and basal layers of the sensory epithelium (SE) (Figure 7). eGFP expression does not localize to either the apical or basal layer, suggesting that eGFP expression occurs in both V1R and V2R subsets of VNO sensory neurons. There is heavy expression of eGFP within the axons of the neurons projecting along the septum posteriorly to the AOB (Figure 7). Interestingly, the septal organ, which lies just posterior to these axonal fibers, is devoid of eGFP expression. The AOB has varied co-expression of eGFP within each glomerulus (Figure 7E), with some glomeruli having a higher intensity of eGFP than others. However, there is no preferential organization between the anterior and posterior AOBs, suggesting that the eGFP-expressing neurons are not segregated based on V1R or V2R expression, consistent with our observations of VSNs.

**Figure 7.**
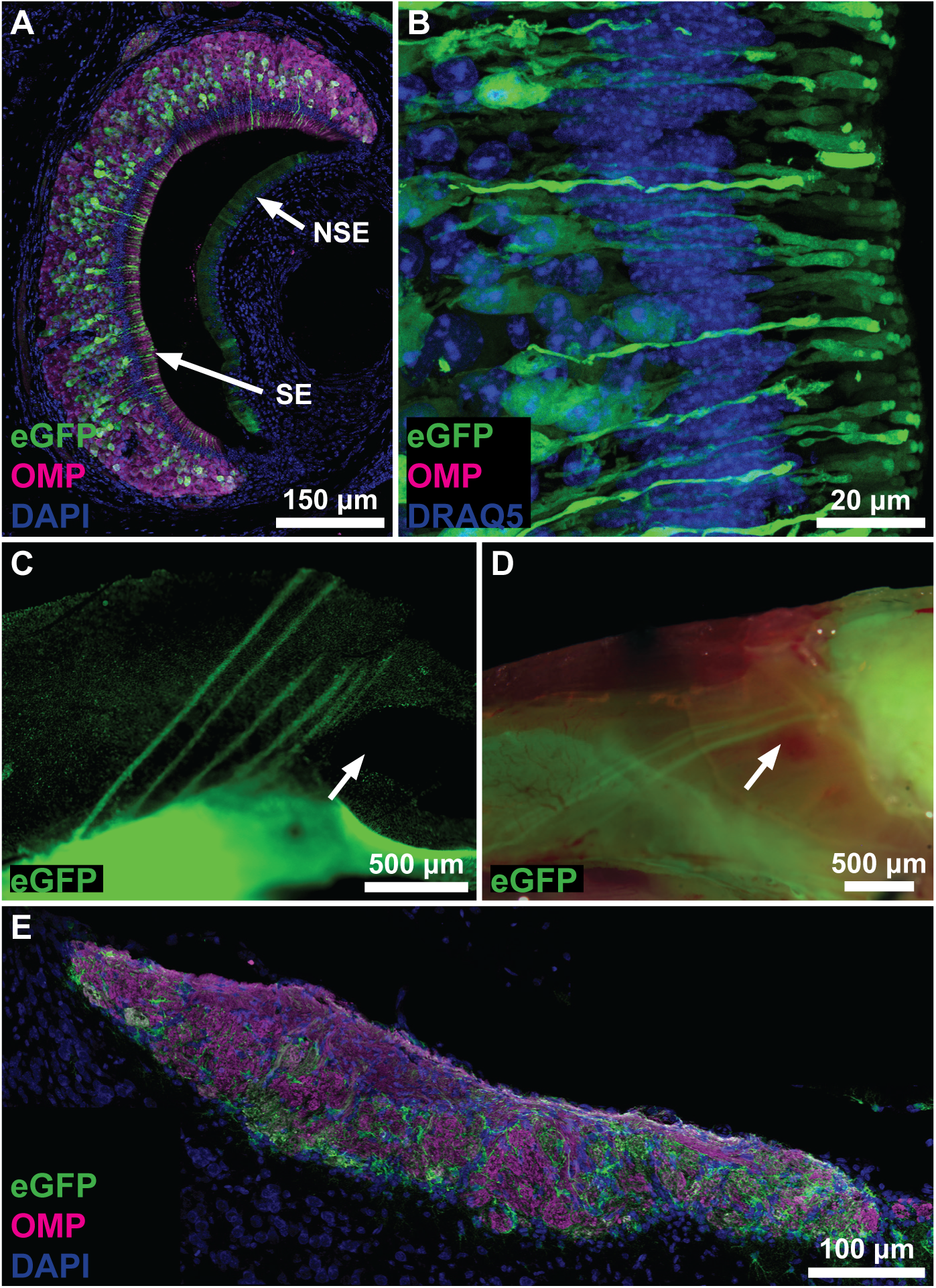
Foxj1-driven eGFP (green) expression in the accessory olfactory system. A. A cross section of the VNO reveals the SE with scattered eGFP expression and the lumen of the VNO. eGFP expression is seen in both the superficial and basal regions of the SE. eGFP-expression also occurs in scattered cells the non-sensory epithelium of the VNO, but at much lower levels relative to the SE. OMP-immunoreactivity; magenta. Nuclear counterstain (DRAQ5); blue. B. High magnification of the apical portion of the SE showing robust eGFP expression throughout the VNO sensory neurons. C, D. Lateral view of two bisected skulls showing eGFP-expressing fibers of the accessory olfactory nerves running from the VNO towards the AOB. In D, the eGFP fluoresence image is overlaid on a color photograph. Septal organ (arrow) is devoid of eGFP expression. Note, the VNO was removed in panel D prior to imaging. E. Longitudinal section through the AOB. Glomeruli in both the anterior and posterior AOB show differential expression of Foxj1-driven eGFP and OMP (magenta) within individual glomeruli.

## Discussion

Here we identify a unique set of olfactory sensory neurons within a transgenic mouse line, in which eGFP expression is linked to the *Foxj1* promoter. These OSNs, which are restricted zonally within the OE, express *Foxj1* driven e-GFP, but no apparent FoxJ1 protein. The GFP-labeled OSNs extend their axons to the MOB and target only a few glomeruli, suggesting that these OSNs express a limited set of odorant receptors. eGFP expression occurs not only in these neurons, but also in the vomeronasal sensory neurons and in astrocytes of the olfactory bulb and forebrain. From the literature(Blatt et al., 1999; Hackett et al., 1995; Lim et al., 1997), we know *Foxj1* to be a driver of motile ciliogenesis and as expected we find robust eGFP in the multiciliated cells of the respiratory epithelium. It is interesting to see eGFP expression in the VSNs and astrocytes, which have no motile cilia. Accordingly, we make no claim that expression of *Foxj1* in these cells is related to ciliogenesis. We do suggest that eGFP expression in this mouse may not be related to *Foxj1* transcription, or that perhaps, *Foxj1* plays an additional role to driving motile ciliogenesis. In all mice examined, we see consistent expression of eGFP within the multiciliated cells of the RE, within a particular and consistent set of OSNs of the OE and thus, we are led to believe that eGFP expression is specific and targeted.

In characterizing these OSNs further, we found that OSNs in the OE had no detectable FOXJ1 protein as seen by immunohistochemistry (Figure 2A). It is possible that *Foxj1* itself may not be transcribed in these neurons and eGFP expression could be driven by other transcription factors. Interestingly, our secondary analysis of single cell RNA sequencing studies have revealed that certain OSNs do in-fact express *Foxj1* mRNA (Fletcher et al., 2017; Hanchate et al., 2015; Tan, Li, & Xie, 2015). These *Foxj1*-expressing OSNs express predominant *Olfr* genes predicted to be expressed in Zones 1 and 2 according to Tan et al. (2018; data not shown), fitting with our observations of zonal restrictions of these OSNs.

We found that the axons of the eGFP expressing ORNs target specific glomeruli, consistent with our hypothesis that these neurons must also express a particular set of odorant receptors. The MOE is classically subdivided into four zones (I-IV) based on receptor expression (Mori et al., 2006; Mori et al., 2000). OR expression is restricted to neurons located within a particular zone. There are approximately 1300 odorant receptor (OR) genes in mice (Zhang & Firestein, 2002). Each mature OSN expresses only one olfactory receptor (OR) and responds to a limited set of odorants (Hanchate et al., 2015). Although cells expressing the same receptor are scattered within one zone in the MOE, their axons converge onto only 1 or 2 glomeruli on each side of the MOB, such that each glomerulus within the MOB receives input from ORNs expressing the same OR (Breer et al., 2006). Our observation that eGFP expression is zonally restricted within the MOE and that they converge onto specific glomeruli within the OB suggest that these neurons express a specific OR. Additionally, the eGFP positive neuronal cell bodies lie at a similar apical distance in regard to their vertical position within the MOE, which also suggests expression of a specific OR. Strotmann et al, (Strotmann et al., 1996) studied laminar segregation of sensory neurons expressing distinct receptors by utilizing *in situ* hybridization employing receptor-specific probes. They found laminar segregation of neurons based on OR expression, such that neurons expressing the same OR were located consistently at a particular depth within the epithelial layer and that its location was unique to the specific OR (Strotmann et al., 1996). Furthermore, in support of the idea that this subpopulation expresses a particular OR, individual glomeruli were identified within the MOB with robust eGFP expression, indicating axonal convergence of these neurons, a pattern seen amongst neurons expressing the same OR.

The location of these eGFP expressing OSNs within an area close to zone 1 or zone 2 of the OE remains of particular interest. The eGFP population resembles the mOR37 family of olfactory receptors described by Breer et. al as they are both restricted to Zone 1 of the OE. Although these neurons are similar in laminar segregation, the mO37 neurons appear to be more restricted within its zone as opposed to the eGFP expressing population that are more scattered (Hoppe, Weimer, Beck, Breer, & Strotmann, 2000).

The expression of the transcription factor *Foxj1* drives motile ciliogenesis in the respiratory epithelium is regulated and driven by upstream multiciliated cell transcription factors including Myb, P73 and those in the E2F and RFX family. (You et al., 2004; Yu et al., 2008). Foxj1 is not required for multiciliated cell fate determination or the initiation of motile ciliogenesis. It plays a role in the apical transport of ciliary basal bodies and the assembly and maintenance of motile axonemes.

Chen et al, (Chen, Knowles, Hebert, & Hackett, 1998) showed that *Hfh4* (*Foxj1*) null mice showed complete absence of cilia and random determination of left-right asymmetry, indicating that *Hfh4* (*Foxj1*) is essential for the development of ciliated cells. The group also found that *Hfh4* (*Foxj1*) may act by regulating expression of the dynein family of genes, which are important in ciliary motility (Chen et al., 1998). The importance of *Foxj1* and motile ciliogenesis was further confirmed *in vitro*, where You et al. showed that cultured *Foxj1* null cells do not develop cilia (You et al., 2004).

Olfactory cilia are similar in structure to the motile cilia found in respiratory epithelium as they both share the 9+2 microtubule configuration (Menco, 1984). OSNs, however, are traditionally categorized as lacking the dynein arms necessary for motility (Menco, 1984). The expression of *Foxj1*, a transcriptional regular known to regulate dynein expression (You et al., 2004), within a subset of these neurons indicate a possibility that these cilia may actually harbor dynein proteins and thus possess motile cilia. We inspected ciliary motility of isolated eGFP expressing OSNs and did not find evidence for ciliary motility (data not shown). Furthermore, cells without cilia, such as the microvillar neurons within the VNO and glial cells within the OB also express FoxJ1-driven GFP, and as such we make no claim that the eGFP expressing OSNs harbor motile cilia.

*Foxj1* - expression also has been characterized recently in neural progenitor cells, including astrocytes and ependymal cells (Jacquet et al., 2011). Expression of *Foxj1* in these cells is thought to be required for embryonic-to-postnatal transition of neurogenesis in the olfactory bulb, perhaps through a paracrine signaling mechanism (Jacquet et al., 2011). Yet, it is still unclear why glial cells of the olfactory bulb exhibit such strong eGFP fluorescence.

The *Foxj1* positive OSN is a new and unique subtype discovered within the olfactory system of mice. The organization pattern seen amongst these neurons suggests that it expresses a limited subset of odorant receptors and characterization of such receptors and corresponding odorants would be an interesting topic of further research.

## Acknowledgements

Special thanks to Dr. Lawrence Ostrowski (UNC Chapel Hill) for generously providing the *Foxj1*-eGFP mouse line, shared by Dr. Eszter Vladar.

